# Thoracolumbar intervertebral disc area morphometry in elderly Chinese men and women: radiographic quantifications at baseline and changes at year-4 follow-up

**DOI:** 10.1101/139402

**Authors:** Jùn-Qīng Wáng, Zoltán Káplár, Min Deng, James F. Griffith, Jason C. S. Leung, Anthony WL Kwok, Timothy Kwok, Ping Chung Leung, Yì-Xiáng J. Wáng

## Abstract

The manuscript submitted does not contain information about medical device(s)/drug(s). No benefits in any form have been or will be received from a commercial party related directly or indirectly to the subject of this manuscript.

**Study Design:** A population-based radiographic study with longitudinal follow-up.

**Objective:** To develop a quantitative index for lumbar disc space narrowing (DSN) evaluation in elderly subjects; to determine how DSN in the elderly is influenced by osteoporosis and gender.

**Summary of Background Data:** There is paucity of research on quantitative classification of lumbar DSN based on disc areal morphometry.

**Methods:** With the database of Osteoporotic Fractures in Men (Hong Kong) and Osteoporotic Fractures in Women (Hong Kong) Studies and those who attended the year-4 follow-up (n = 1519 for men and n = 1546 for women), data of 491 women and 592 men were randomly selected. The anterior, middle, and posterior heights, anteroposterior diameter and area of intervertebral discs (T4T5 to L4L5) were measured on lateral radiographs. Disc Area Index for Lumbar Spine (DAIL, disc area divided by the mean of the sum of square of the adjacent upper and lower vertebrae mid-height anterior-posterior diameter) was developed and compared with semi-quantitative DSN expert grading.

**Results:** DAIL correlated with semi-quantitative grading, with sensitivity and specificity varying from 87.3% to 96.8% for grade-1 DSN (<30% reduction in disc height), and 92.9 % to 100% for grade-3 DSN (>60% reduction in disc height). The thoracolumbar disc area loss among men and women during 4-years’ follow-up period varied between 1.32% and 3.56%, and it was greater for women (mean: 2.44%) than for men (mean: 1.90%, *p*=0.044). Majority of lumbar DSN progressions during 72 to 76 years old were progression from normal disc space to grade-1DSN. Osteoporosis was associated with greater disc area decrease, both for thoracic and lumbar discs.

**Conclusion:** Lumbar DSN can be quantified using DAIL. In elderly Chinese, intervertebral disc narrowing over a 4-year period was greater in women than men, and associated with the presence of osteoporosis.

## Introduction

Spine degeneration is commonly associated with osteophytes formation, decreased bone mineral density (BMD), decrease of vertebral body middle height (i.e. increased biconcavity), increased wedge of thoracic vertebral bodies, and osteoporotic fracture. Intervertebral disc degeneration can progress to disc herniation, spinal canal stenosis, and, in conjunction with facet joint arthrosis, degenerative spondylolisthesis [1–5]. Histology studies show disc degeneration becomes apparent in men in the second decade of life, almost a decade earlier than in women [6, 7]. While young and middle-aged men are more likely to have lumbar disc degeneration than women, radiological evidences demonstrate this trend is reversed in elderly subjects, with women tending to have more severe lumbar disc degeneration than men [8, 9], and this lead to increased low back pain incidence in postmenopausal women compared with age-match men [10]. There are evidences to suggest that osteoporosis, disc degeneration (loss of disc height), and spine fracture interplay with each other. For example, disc degeneration transfers load bearing from the anterior vertebral body to the neural arch in upright postures, reduces BMD and trabecular architecture anteriorly, and predisposes vertebral body to anterior fracture when the spine is flexed [11]. Osteoporotic endplate micro-fractures and compromised healing can negatively impact disc nutrition and contribute to disc degeneration [12, 13]. Recently evidences also show that discs and vertebrae degenerate or remodel in concert [14].

Till now the areal loss of thoracic and lumbar disc space and their association with BMD in elderly subjects, and their gender differences, over a defined time span remain unknown. Osteoporotic Fractures in Men (Mr. OS) (Hong Kong) and Osteoporotic Fractures in Women (Ms OS) (Hong Kong) represent the first large-scale prospective cohort studies ever conducted on bone health in Asian men and women. Utilizing this database, the purpose of the current study was threefolds: 1) Till now, the diagnose of intervertebral disc space narrowing is subjective and uses a semi-quantitative grading, we aim to develop a quantitative index for lumbar disc space narrowing evaluation in elderly subjects; 2) to quantify the areal loss of thoracic and lumbar disc space over four years in elderly females and males; 3) to further confirm the previous observation that osteoporosis is associated with faster disc volume loss than normal BMD subjects [15].

## Materials and methods

Mr. OS (Hong Kong) and Ms OS (Hong Kong) studies design follow that of the osteoporotic fracture in men (MrOS) study performed in the United States [16]. At baseline, 2,000 Chinese men (mean age: 72.39 yrs) and 2,000 Chinese women (mean age: 72.58 yrs) in Hong Kong aged 65 to 98 years were recruited from the local communities between August 2001 and March 2003 [17, 18]. The recruitment criteria were established so that the study results from the cohort would be applicable to a broad population of similarly aged community-dwelling men and women. The project was designed primarily to examine the bone mineral density (BMD) of older Chinese adults prospectively for 4 years. All participants were community dwelling, able to walk without assistance, had no bilateral hip replacement and had the potential to survive the duration of a primary study based on their general medical health. The study protocol was approved by the Chinese University of Hong Kong Ethics Committee. 1,519 males (76.0%) and 1,546 females (77.3%) attended the year-4 follow-up study [19]. The remaining participants were unwilling or unable to attend for follow-up or were not contactable.

BMD (g/cm^2^) at the total hip was measured by Hologic QDR 4,500 W densitometers (Hologic Inc., Waltham, MA). Subjects were divided into three groups, i.e., normal BMD, osteopenia, and osteoporosis, according to World Health Organization criteria. A subject is defined as being normal if their T-score is above −1.0; osteopenic if their T-score is between −1.0 and −2.5; and osteoporotic if their T-score is below −2.5 [20]. Standard Hong Kong Chinese reference data were used for the T-score calculations [18, 21]. Spine radiographs were centered on T7 for the thoracic spine (T3-L1) and on L3 for the lumbar spine (T12-S1). Left lateral thoracic and lumbar spine radiographs were obtained by adjusting exposure parameters according to participants’ body weight and height. The standard parameters were: thoracic spine: -Film/Focus Distance: 40 inches, voltage 60-70 kVp, Exposure Time: 2 seconds; and lumbar spine: -Film/Focus Distance: 40 inches -Imaging voltage 80-90 kVp -Exposure Time: 1sec. These radiograph parameters were the same for baseline and for follow-up. Radiographs were digitized with spatial resolution of 300 dpi using VIDAR’s DiagnosticPRO(r) Advantage film digitizer, and ClinicalExpress(r) 4.0 software (Vidar Systems Corporation, Herndon, USA).

500 women and 600 men’s data were randomly selected from those who attended both baseline and follow-up studies (Fig 1). This sample size estimation was based on previous quantitative MRI study of lumbar vertebrae and lumbar disc [15], and the consideration that thoracic spine discs have smaller size and more difficult to be measured reliably than lumbar discs, and elderly men demonstrates less extent of changes than elderly women with fewer of them having osteoporosis. Data from eight men and nine women were excluded due to inferior radiograph quality. Morphometric measurement was performed in each vertebra from T4 to L5 using a program written with Matlab (Matlab R2015a, Mathworks, USA). Eight digitized reference points were manually placed for each vertebra (Figure 2A), and disc dimensions including anterior height (Ha), middle height (Hm), posterior height (Hp), anteroposterior diameter (AP) and disc areas from T4 to L5 were generated. The disc area was calculated as a hexagonal area composed of 4 triangles, formed by 6 intersecting lines (Figure 2B). For the correction of potential magnification differences between baseline and follow-up radiographs of the same participant, the coordinates of the points from follow-up radiographs was normalized with mid-height AP diameter of vertebral bodies at baseline. Based on past publications [22–24], the assumption was taken that vertebral mid-height AP diameter would not notably change during the 4-yrs follow-up. Similar to previous reports, disc space at L5S1 was not included, as assessment of disc narrowing at this level is less reliable [17, 25]. Under the close supervision of an experienced radiologist (YXJW), two readers performed the morphometric measurement, Reader-1 (JQW) measured the radiographs of 491 females and 250 males, and reader-2 (ZK) measured the remaining 342 males. 50 randomly selected radiographs were measured for reproducibility assessment. The intraclass correlation coefficient (ICC) for intra-reader repeatability was 0.988 (Ha), 0.986 (Hm), 0.979 (Hp), and 0.990 (disc area), respectively; while ICC for inter-observer repeatability was 0.950 (Ha), 0.942 (Hm), 0.922 (Hp) and 0.985 (disc area), respectively.

**Figure 1.**
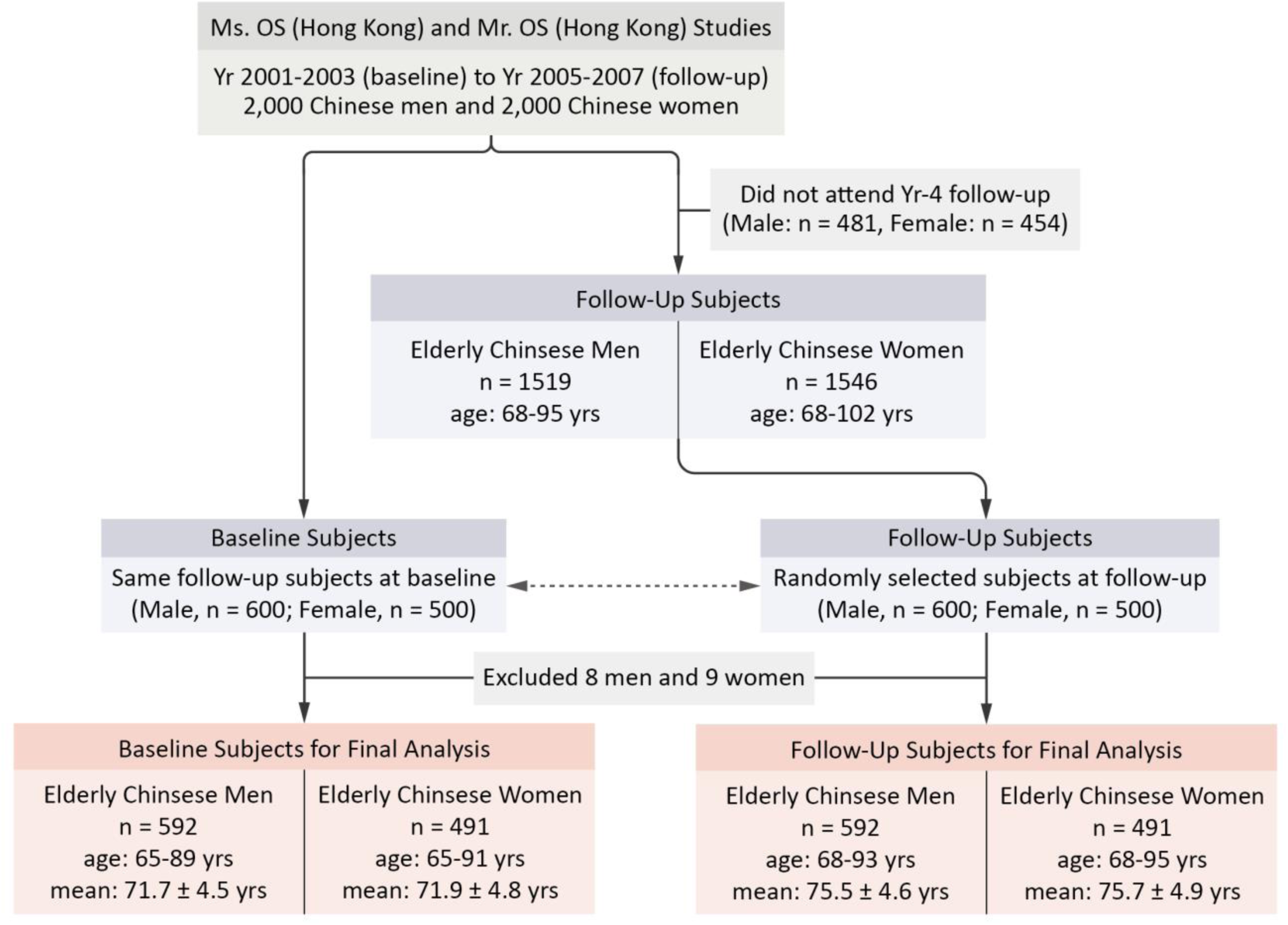
The flow chart shows the selection of study subjects.

**Figure 2.**
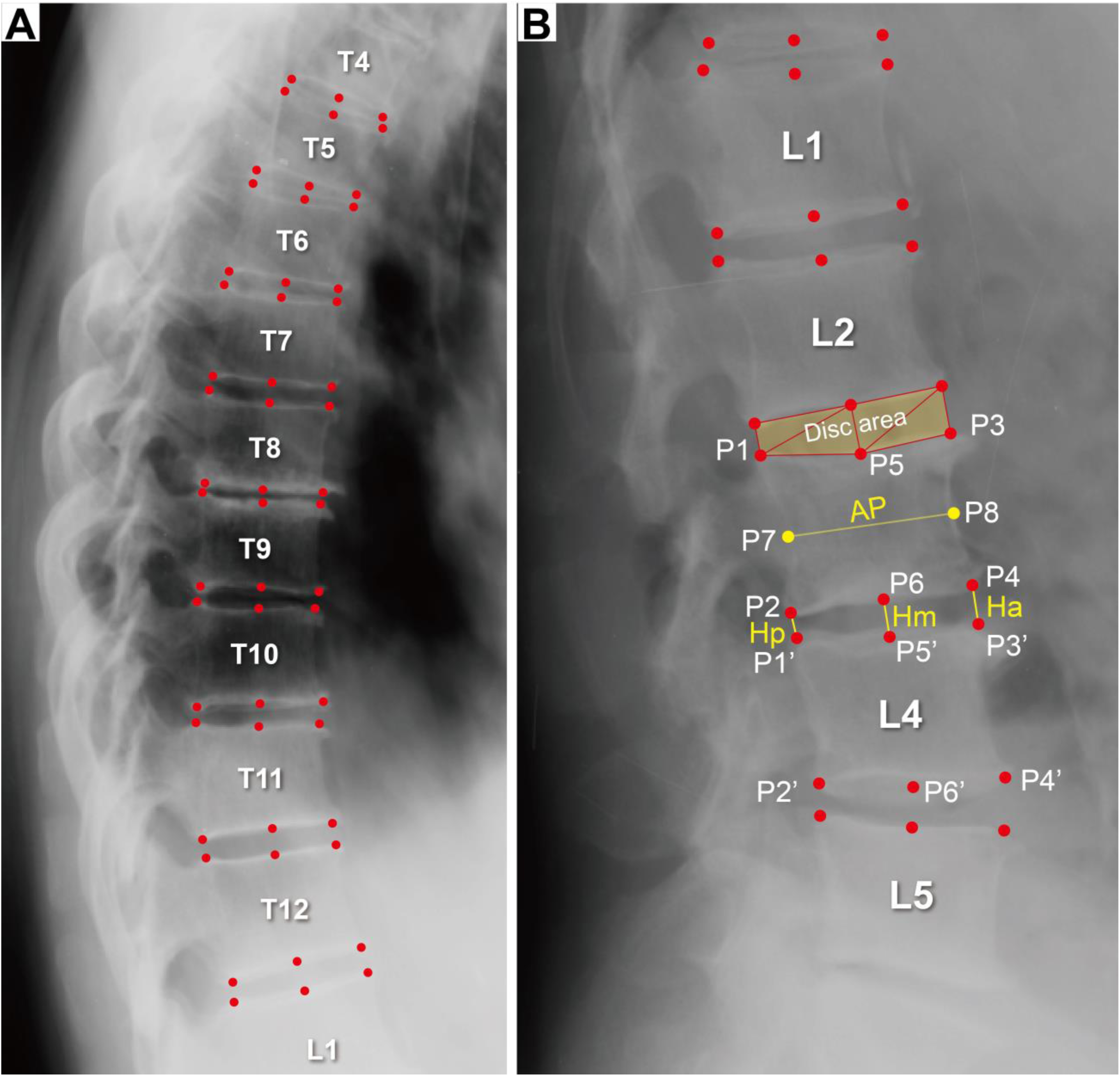
8-point vertebral body and disc morphometry of spinal radiograph. Four contour points (P1-P4) were identified at the four corners of the vertebral body, two midpoints (P5 and P6) were marked at middle of the upper and lower endplates, and additional two points (P7 and P8) were positioned on the middle of the ventral (P1-P2) and dorsal (P3-P4) lines (A). The disc area is presented as a hexagonal area composed of 4 triangles (B). When scoliosis exists and the endplate shows double-lines, points P5 and P6 are placed at the middle points of the two double-lines.

Disc Area Index for Lumbar spine (DAIL) for each intervertebral level at baseline were calculated using the Equation (1, supplementary Fig 1).

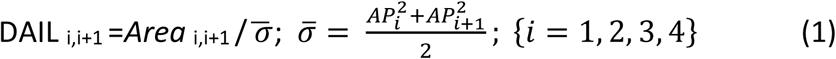

Where *Area* is the intervertebral disc area, *i* = 1, 2, 3, 4 is the vertebral level, *AP* is the mid-height anteroposterior diameter of vertebral body (*AP_i_*: the vertebral body above the disc, *AP*_*i+1*_: the vertebral body below the disc), 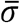 is the mean of the sum of square of the adjacent upper and lower vertebrae anteroposterior diameter *(AP_i_* and *AP*_*i+1*_). Therefore, DAIL refers to the area of a disc divided by an area formed by mid-height anteroposterior diameters of the two adjacent vertebral bodies, and thus is unitless. As the mid-height anteroposterior diameters of the two adjacent vertebral bodies are usually unaffected by spine degeneration, and the narrower the disc space, the smaller the DAIL value. The reference standard grading was from a previous study with this dataset [17]. By experienced radiologists, lumbar disc space was visually classified into 4 categories with the aid of direct measurement for borderline cases: normal (grade-0), mild narrowing (grade-1< 30% reduction in disc height), moderate narrowing (grade-2=30–60% reduction in disc height), and severe narrowing (grade-3>60% reduction in disc height) [17, 25]. DAIL threshold criteria for defining severity of DSN from grade-1 to grade-2 and grade-3 were obtained from receiver operating characteristic (ROC) analysis (Fig 3, Supplementary Fig 3–4). Using these DAIL cut-off values, the lumbar spine radiographs obtained at year-4 follow-up were used to evaluate DSN progression, and then the results were confirmed by a radiologist (MD) who participated in the previous study [17].

**Figure 3.**
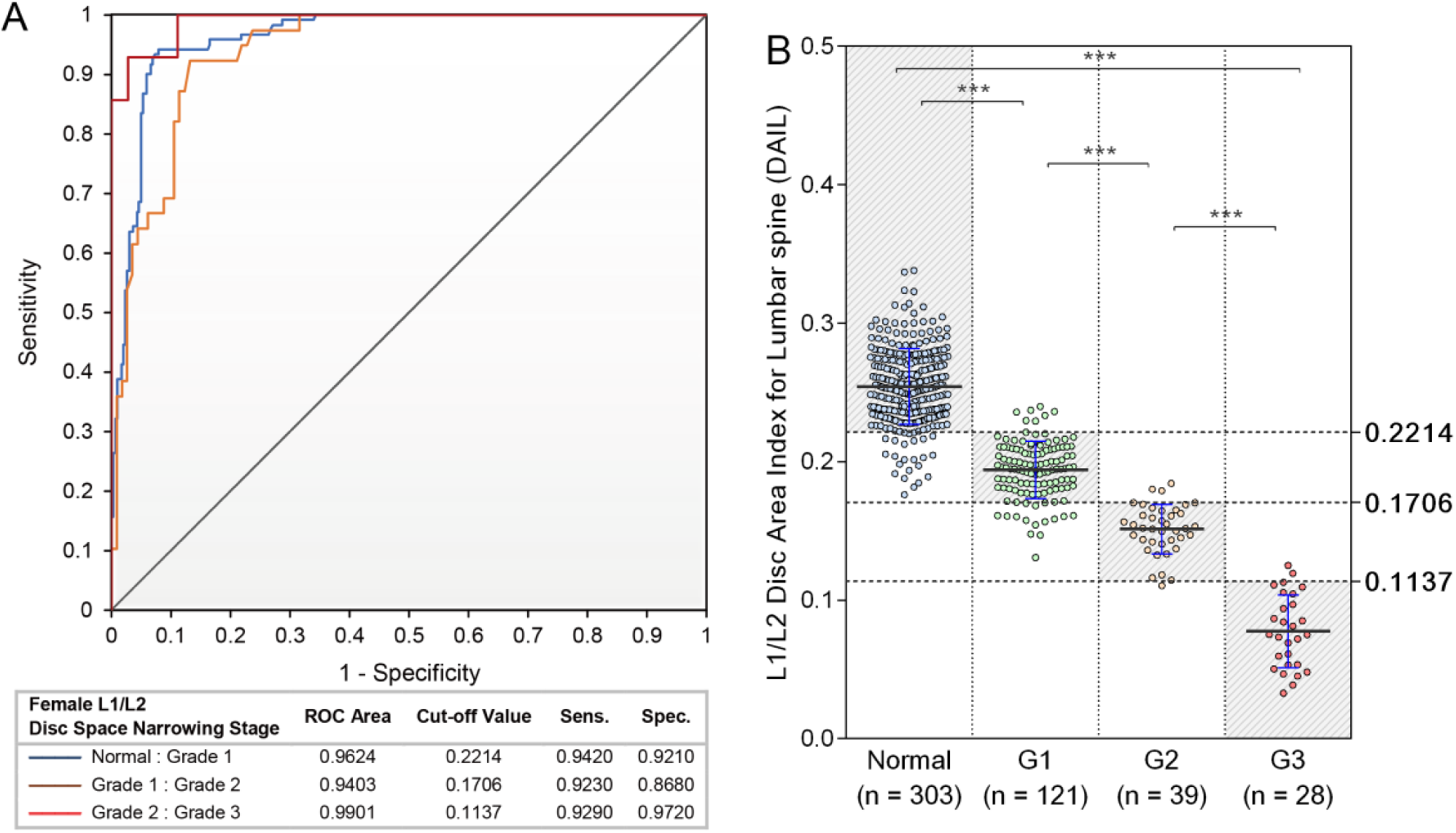
A: an example of ROC curve and diagnostic ability for L1L2 DSN in women; B: scatter plot of L1L2 DAILs which correlate with normal disc space, grade 1, 2 and 3 DSN. Defined optimal cut-off DAILs for DSN grading are indicated by horizontal dash line (more examples see supplementary Figures).

The statistical package IBM SPSS Statistics, V21.0 (IBM Corporation, IBM Corp, Armonk, New York, USA) was used for data processing. A probability level of 0.05 was used as the level of significance.

## Results

The demographic variables of study subjects are summarized in Table 1. There was no difference in age among the male and female groups, and there were more female subjects with osteoporosis than males (18.74% vs 3.72% at baseline, 24.24% vs 3.89% at year-4 follow-up).

**TABLE 1.** Demographics of Study Subjects

|  | Women (N = 491) |  | Men (N = 592) |  |
| --- | --- | --- | --- | --- |
|  | BL | FU | BL | FU |
| Mean age (yrs) ± SD (range) # | 71.9 ± 4.8 (65-91) | 75.7 ± 4.9 (68-95) | 71.7 ± 4.5 (65-89) | 75.5 ± 4.6 (68-93) |
| Mean height (cm) ± SD | 151.7 ± 5.2 | 151.1 ± 5.3 | 163.2 ± 5.5 | 162.8 ± 5.5 |
| Mean weight (kg) ± SD | 55.3 ± 8.3 | 54.5 ± 8.6 | 63.3 ± 8.8 | 62.5 ± 8.7 |
| Normal BMD subjects | 135/491 (27.49%) | 120/491 (24.44%) | 311/592 (52.53%) | 297/592 (50.17%) |
| Osteopenia subjects | 264/491 (53.77%) | 252/491 (51.32%) | 259/592 (43.75%) | 272/592 (45.95%) |
| Osteoporosis subjects | 92/491 (18.74%) | 119/491 (24.24%) | 22/592 (3.72%) | 23/592 (3.89%) |
| BL: baseline; FU: year-4 follow-up; # *p* for women vs men at BL=0.508 |  |  |  |  |

The ROC analysis determined DAIL cut-off criteria for classifying lumbar DSN from grade-1 to grade-3 are shown in Table 2 and Supplementary Figures 3-4. DAIL correlated well with semi-quantitative grading, with sensitivity and specificity varying from 87.3% to 96.8% for grade-1 DSN, and 92.9 % to 100% for grade-3 DSN. DAIL performed the best at grade-3 DSN, and the performance was slightly lower for grade-1 DSN.

**TABLE 2.** Receiver operating characteristic (ROC) analysis of DAIL-based DSN Classification for lumbar discs at Baseline

|  | L1L2 |  |  | L2L3 |  |  | L3L4 |  |  | L4L5 |  |  |
| --- | --- | --- | --- | --- | --- | --- | --- | --- | --- | --- | --- | --- |
| Females | G1 | G2 | G3 | G1 | G2 | G3 | G1 | G2 | G3 | G1 | G2 | G3 |
| AUC | 0.96 | 0.94 | 0.99 | 0.94 | 0.97 | 0.99 | 0.96 | 0.97 | 1.00 | 0.98 | 0.97 | 0.98 |
| 95% CI | 0.946-0.979 | 0.910-0.977 | 0.976-1.00 | 0.919-0.967 | 0.955-0.994 | 0.985-1.000 | 0.947-0.980 | 0.960-0.990 | 0.994-1.000 | 0.964-0.992 | 0.951-0.984 | 0.951-1.000 |
| p | <0.0001 | <0.0001 | <0.0001 | <0.0001 | <0.0001 | <0.0001 | <0.0001 | <0.0001 | <0.0001 | <0.0001 | <0.0001 | <0.0001 |
| Sensitivity (%) | 94.2 | 92.3 | 92.9 | 87.3 | 92.5 | 94.9 | 93.8 | 95.1 | 100.0 | 96.8 | 92.3 | 94.9 |
| Specificity (%) | 92.1 | 86.8 | 97.2 | 94.2 | 95.2 | 98.0 | 88.6 | 90.5 | 97.4 | 94.3 | 89.0 | 97.7 |
| DAIL cut-off value | 0.2214 | 0.1706 | 0.1137 | 0.2378 | 0.1787 | 0.1147 | 0.2625 | 0.2025 | 0.1208 | 0.3008 | 0.2244 | 0.1253 |
| Males | G1 | G2 | G3 | G1 | G2 | G3 | G1 | G2 | G3 | G1 | G2 | G3 |
| AUC | 0.96 | 0.98 | 1.00 | 0.96 | 0.98 | 0.99 | 0.97 | 0.99 | 1.00 | 0.98 | 0.98 | 1.00 |
| 95% CI | 0.950-0.979 | 0.962-0.999 | 1.000-1.000 | 0.937-0.976 | 0.971-0.998 | 0.976-1.000 | 0.954-0.984 | 0.984-0.999 | 0.995-1.000 | 0.966-0.991 | 0.967-0.992 | 0.989-1.000 |
| p | <0.0001 | <0.0001 | <0.0001 | <0.0001 | <0.0001 | <0.0001 | <0.0001 | <0.0001 | <0.0001 | <0.0001 | <0.0001 | <0.0001 |
| Sensitivity (%) | 92.3 | 92.9 | 100.0 | 90.5 | 96.4 | 92.9 | 92.3 | 95.5 | 100.0 | 95.2 | 92.0 | 97.9 |
| Specificity (%) | 91.2 | 96.4 | 100.0 | 91.0 | 92.6 | 100.0 | 95.6 | 97.1 | 97.0 | 92.7 | 95.2 | 96.9 |
| DAIL cut-off value | 0.2345 | 0.1775 | 0.1157 | 0.2662 | 0.1995 | 0.1208 | 0.2784 | 0.2185 | 0.1552 | 0.3194 | 0.2484 | 0.1574 |

DSN: disc space narrowing; AUC: area under the curve; G1 G2, G3: DSN of grade 1, 2, and 3, respectively.

At the year-4 follow-up, the agreement between DAIL-based and radiologist DSN gradings had a kappa value of 0.745 for women, and 0.732 for men. The progression of lumbar DSN during 72 to 76 years were mostly from normal to grade-1 (Table 3). In females the proportion of normal spaced discs decreased from 45.1% at baseline to 36.6% at year-4 follow-up, while in males the proportion of normal spaced discs decreased from 49.2% to 40.8%.

**TABLE 3.** Progress of Lumbar Disc Space Narrowing During the 4-Years Follow-Up Period for Women and for Men (based on DAIL read)

|  |  |  |  |  |  |
| --- | --- | --- | --- | --- | --- |
| Disc Level | DSN classification | Women Baseline # | Women Yr-4 # | Men Baseline# | Men Yr-4 # |
| Total (L1L2 – L4L5) | Grade 1 | 30.40% | 38.14% | 33.61% | 38.72% |
| Total (L1L2 – L4L5) | Grade 2 | 17.57% | 17.92% | 13.22% | 14.44% |
| Total (L1L2 – L4L5) | Grade 3 | 6.93% | 7.38% | 3.93% | 6.04% |
| L1L2 | Grade 1 | 25.66% | 41.96% | 28.55% | 35.64% |
|  | Grade 2 | 10.59% | 12.22% | 4.73% | 6.93% |
|  | Grade 3 | 5.50% | 6.31% | 1.52% | 2.53% |
| L2L3 | Grade 1 | 27.90% | 37.07% | 31.93% | 38.34% |
|  | Grade 2 | 11.41% | 12.02% | 9.46% | 12.50% |
|  | Grade 3 | 7.74% | 8.15% | 2.36% | 3.38% |
| L3L4 | Grade 1 | 31.57% | 35.85% | 35.14% | 39.53% |
|  | Grade 2 | 17.92% | 18.53% | 11.15% | 12.50% |
|  | Grade 3 | 6.11% | 6.31% | 3.72% | 7.26% |
| L4L5 | Grade 1 | 36.46% | 37.68% | 38.85% | 41.39% |
|  | Grade 2 | 30.35% | 28.92% | 27.53% | 25.84% |
|  | Grade 3 | 8.35% | 8.76% | 8.11% | 10.98% |

DSN: disc space narrowing. #: the portions of total male/female subjects classified with a specific DSN grade.

Thoracic and lumbar lateral disc area decreases during 4-years follow-up period are shown in table 4 (supplementary table 1 and 2). There was a statistically significant trend that lower hip BMD measured at baseline year was associated with greater disc area loss during the 4-year period. The thoraco-lumbar disc area losses among men and women during 4 years’ follow-up period varied between 1.32% and 3.56%, and it was greater for women (mean: 2.44%) than for men (mean: 1.90%, *p* = 0.044, Fig 4). An overall trend was noted that caudal discs had higher percentage area decrease than cephalad discs. Both for females and males, in the thoracic spine there was a greater percentage disc area loss in mid-thoracic region than lower thoracic region (Fig 4).

**TABLE 4.** Female and Male Lateral Intervertebral Disc Area loss in 4-Years among normal BMD, Osteopenia, and Osteoporosis subjects

|  |  |  |
| --- | --- | --- |
|  | Estimated Means of Disc Area loss in 4 years* |  |
| Female | Thoracic Discs (mean±SD) | Lumbar Discs (mean±SD) |
| Total (n = 491) | 1.74% ± 0.058 | 3.56% ± 0.046 |
| Normal BMD (n = 135) | 1.23% (0.005) | 2.51% (0.004) |
| Osteopenia (n = 264) | 1.52% (0.004) | 3.67% (0.003) |
| Osteoporosis (n = 92) | 3.10% (0.007) | 4.79% (0.005) |
| p in linear trend | 0.037 | 0.001 |
| Male |  |  |
| Total (n = 592) | 1.32% ± 0.066 | 2.84% ± 0.042 |
| Normal BMD (n = 311) | 0.85% (0.004) | 2.30% (0.002) |
| Osteopenia (n = 259) | 1.66% (0.004) | 3.34% (0.003) |
| Osteoporosis (n = 22) | 3.89% (0.014) | 4.42% (0.009) |
| p in linear trend | 0.042 | 0.026 |

\* Disc Area loss in 4 years = [(baseline area- follow-up area)/baseline area] × 100%, with analysis of covariance (ANCOVA) and adjustment of BMI (body mass index) and age at baseline.

**Figure 4.**
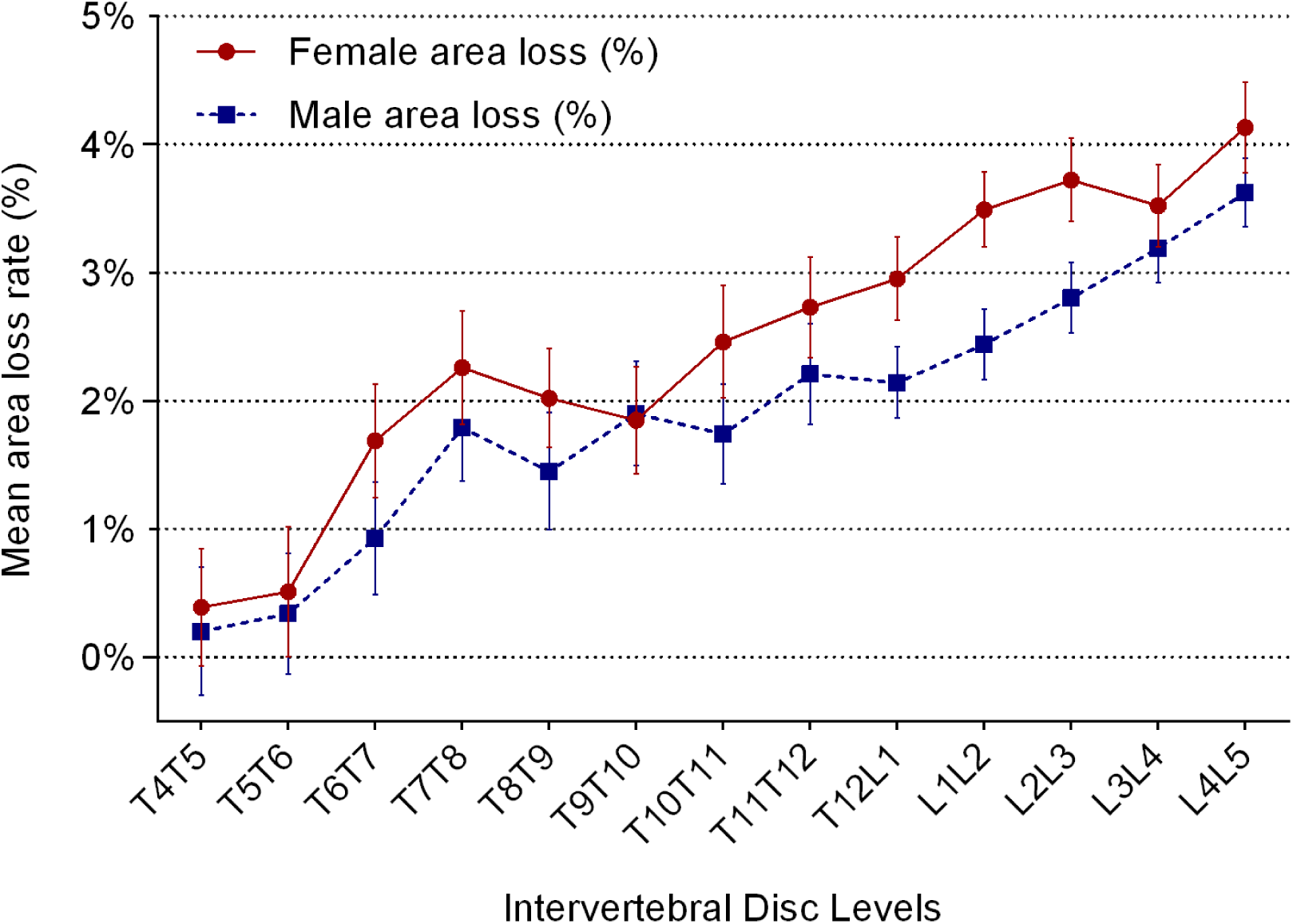
Percentage lateral disc area decrease (mean ± standard deviation) at individual levels during 4-years follow-up, calculated by [(disc area at baseline-disc area at follow-up)/ disc area at baseline period]. Female subjects have a higher lateral disc area loss rate than males at each disc levels.

## Discussion

This study is the first to investigate the influence of ageing and osteoporosis on the morphology of both thoracic and lumbar intervertebral discs, using quantitative radiographic data for both genders selected from an elderly population at baseline and at year-4 follow-up. One strength of this study is that men and women of similar age and from the same community-based population were investigated, thereby enabling men and women to be directly compared.

DSN has been traditionally semi-quantitatively graded by experienced radiologists/physicians [13, 16]. However, such semi-quantitative grading is subjective, making it difficult for epidemiological study and longitudinal follow-up. Our study developed DAIL, which can quantitatively classify lumbar disc space into normal and DSN. The DAIL criteria was tested to compute the DSN progression at year-4 follow-up, and showed good agreement between results of DAIL-based reads and radiologist-based reads, with an overall kappa value of 0.745 for women, and 0.732 for men. These kappa values are similar to the inter-reader reproducibility of a kappa value of 0.72 by two experienced radiologists, which was obtained using the baseline L1/L2-L4/L5 radiographs [17]. In addition to the mild/moderate/severe DSN criteria used in this study, other cut-off values have been proposed. For example, Mimura *et al* (1994) proposed normal, and mild (>75%), moderate (>50%), and severe (>25%), and very severe (<25%) DNS [26]. It should be noted the DAIL cut-off criteria can be re-adjusted to meet these criteria. Computer-aided segmentations for both vertebral body and disc area on lateral radiograph have been developed [27–30]. It is expected that this DAIL criteria method will aid in computerized disc segmentation and automatic DSN grading. On the other hand, visual radiological assessment of radiographs can derive additional information such as discrete endplate defects (e.g. Schmorl’s nodes), the presence of marginal vertebral body osteophytes, etc., all of which give valuable additional information concerning degenerative changes of the spine. Therefore DAIL cannot replace expert evaluation of lumbar radiographs.

Recently evidences suggest relative estrogen deficiency may contribute to the accelerated disc degeneration seen in postmenopausal women [8; 9; 31; 32], which in turn is associated increased prevalence of lower back pain [10]. The current study showed during the 4-years follow-up period there was greater lateral disc area loss in females, and during the period there were more DSN grade progresses in women than in men. This result differs from the report of Gambacciani *et al* [28]. Gambacciani *et al* reported after menopause disc space shows a progressive decrease that almost entirely occurs in the first 5–10 years since menopause. The results of this study, i.e. females have faster disc space narrowing than male even 20 years after menopause, concur with previous reports of Wang *et al* [17] and De Schepper EI *et al* [25]. Our results also showed the lumbar DSN progression mainly occurred from normal disc space to grade-1 DSN in both genders during the follow-up period (7.7 % for women, 5.1 % for men).

A trend was noted that caudal discs had higher lateral area decrease rate than cephalad discs (Fig 4). It has been previously recognized that lumbar discs are more likely to undergo disc degeneration than thoracic discs [33], lower lumbar discs are more likely to undergo severe degeneration than upper lumbar discs [34]. Interestingly, both for females and males, in the thoracic spine there was greater disc area loss in mid-thoracic region than lower thoracic region. This result may be associated with curvature of the spine. The parts with greater spine curvature, i.e. mid-thoracic region and L4/L5, tend to loss lateral disc area more than parts with less spine curvature. Adams *et al* [35] suggested that there are two types of disc degeneration. ‘Endplate-driven’ disc degeneration involves endplate defects and inwards collapse of the annulus, mostly affects discs in the upper lumbar and thoracic spine, usually is associated with compressive injuries. ‘Annulus-driven’ disc degeneration involves a radial fissure and/or a disc prolapse, mostly affects discs in the lower lumbar spine, and is associated with repetitive bending and lifting. Lower lumbar discs are subjected to greater loading in bending, and so are more susceptible to degenerative changes (including disc prolapse) which arise from bending injuries to the annulus. Mid-thoracic discs are more likely to sustain compression injury to an endplate. Therefore, the results of this study may support the observation that two types of degeneration phenotype exist [35].

A trend was significant for a lower baseline BMD associated with a greater decrease of lateral disc areas, both for thoracic and lumber discs among females and males. Previous volumetric MR data suggested that although lower BMD is associated with greater disc middle height and increased biconvexity, lower BMD is accompanied by a decrease in disc volume [15]. Osteoporosis can cause endplate thinning and micro-fracture which in turn lead to compromised endplate healing, and add calcification and decrease the vascularization in the endplates adjacent to the degenerated discs, which subsequently exacerbated degeneration of the associated discs 13, 36-38]. It is noted that for osteoporotic subjects in this study, elderly men and elderly women had similar extent of disc area loss during the 4-years follow-up (table 4).

There are a number of limitations of this study. The DAIL criteria was validated at year-4 follow-up and compared with radiologist reads. However, radiologist DSN grading is itself subjective and could not be considered as golden standard. The DAIL criteria was only validated in elderly Chinese population, how it should be adjusted in younger population or other ethnic groups remain to be further studied. The year-4 follow-up quantification was based on the assumption that there was no change in vertebral mid-height horizontal AP diameter. Though this is a reasonable consideration for the 4-years follow-up period, this may not be absolutely true for individual cases.

In conclusion, the DAIL proposed in this study has a good performance in identifying DSN and may help to standardize automatic grading. In elderly Chinese, intervertebral disc narrowing over a 4 year period was greater in women than men, and associated with the presence of osteoporosis, and was greatest in the lower lumbar spine.

---

**Supplement Figure 1.**
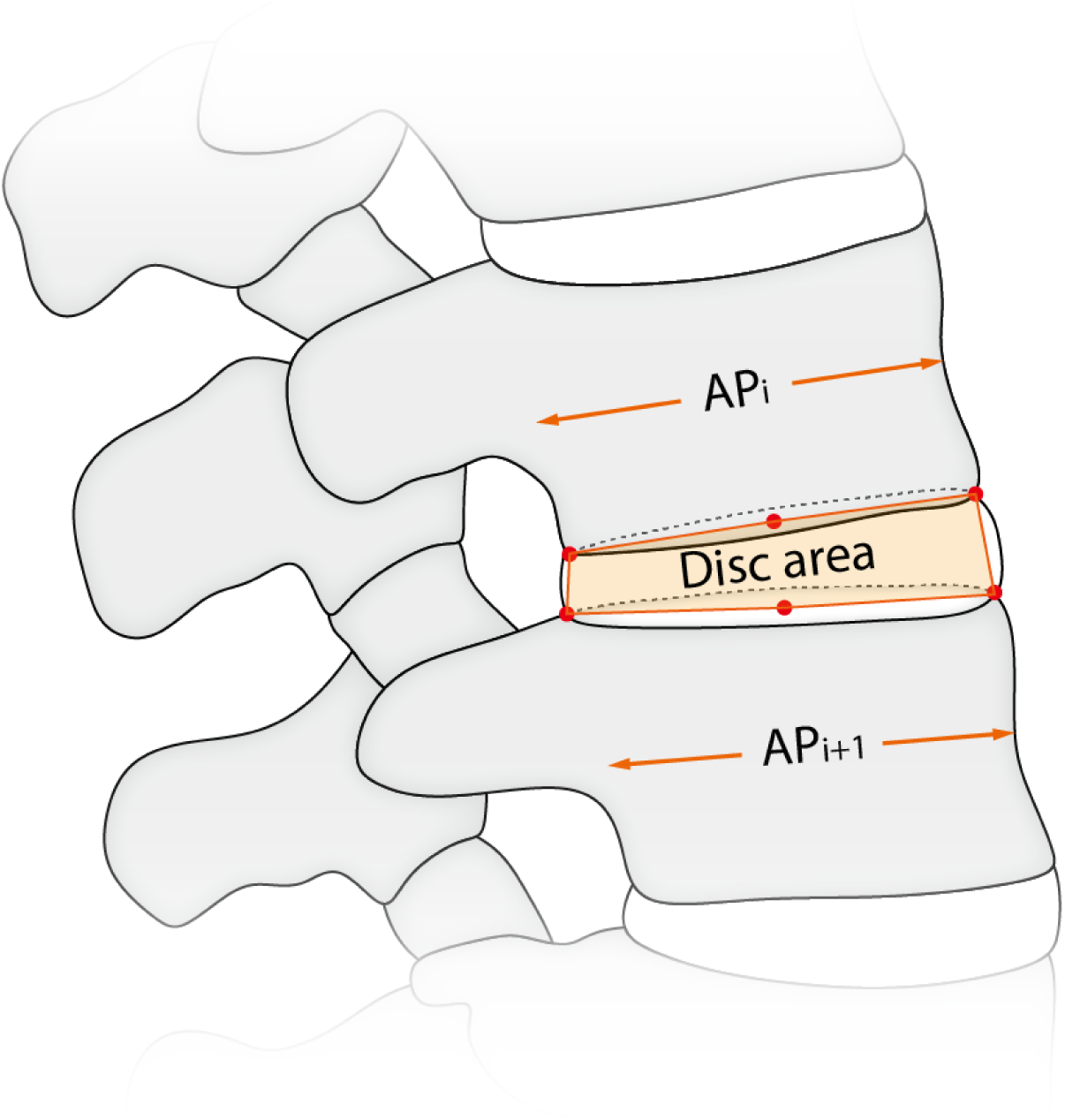
Illustration of relationship of disc area vs. mid-height anterior-posterior (AP) diameter of two adjacent vertebral bodies.

**Supplement Figure 2.**
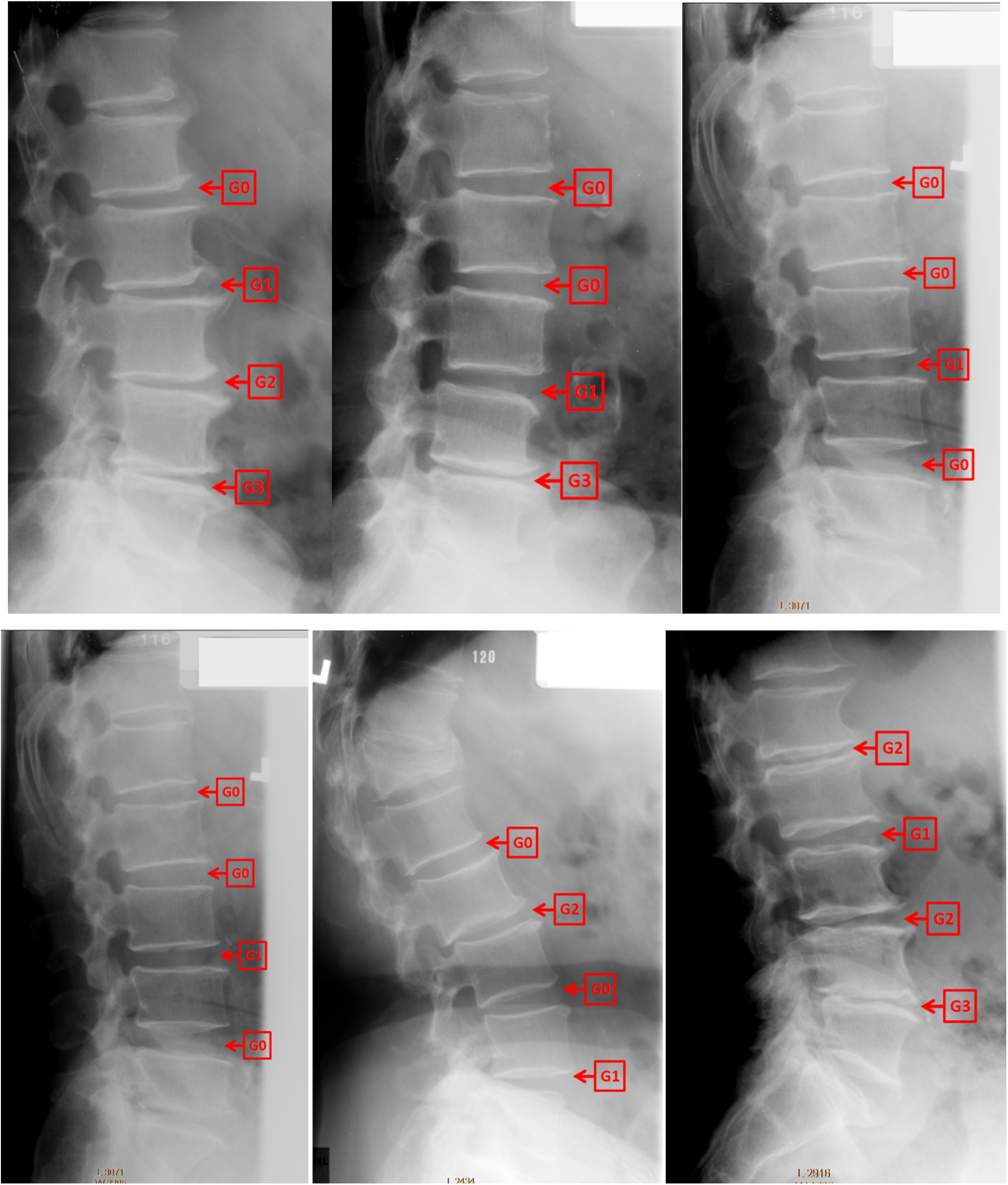
Illustrative examples of visual grading of disc space narrowing. Lumbar disc space is classified into 4 categories: normal (grade 0), mild narrowing (grade-1 < 30% reduction in disc height), moderate narrowing (grade-2 = 30–60% reduction in disc height), and severe narrowing (grade-3 > 60% reduction in disc height).

**Supplement Figure 3.**
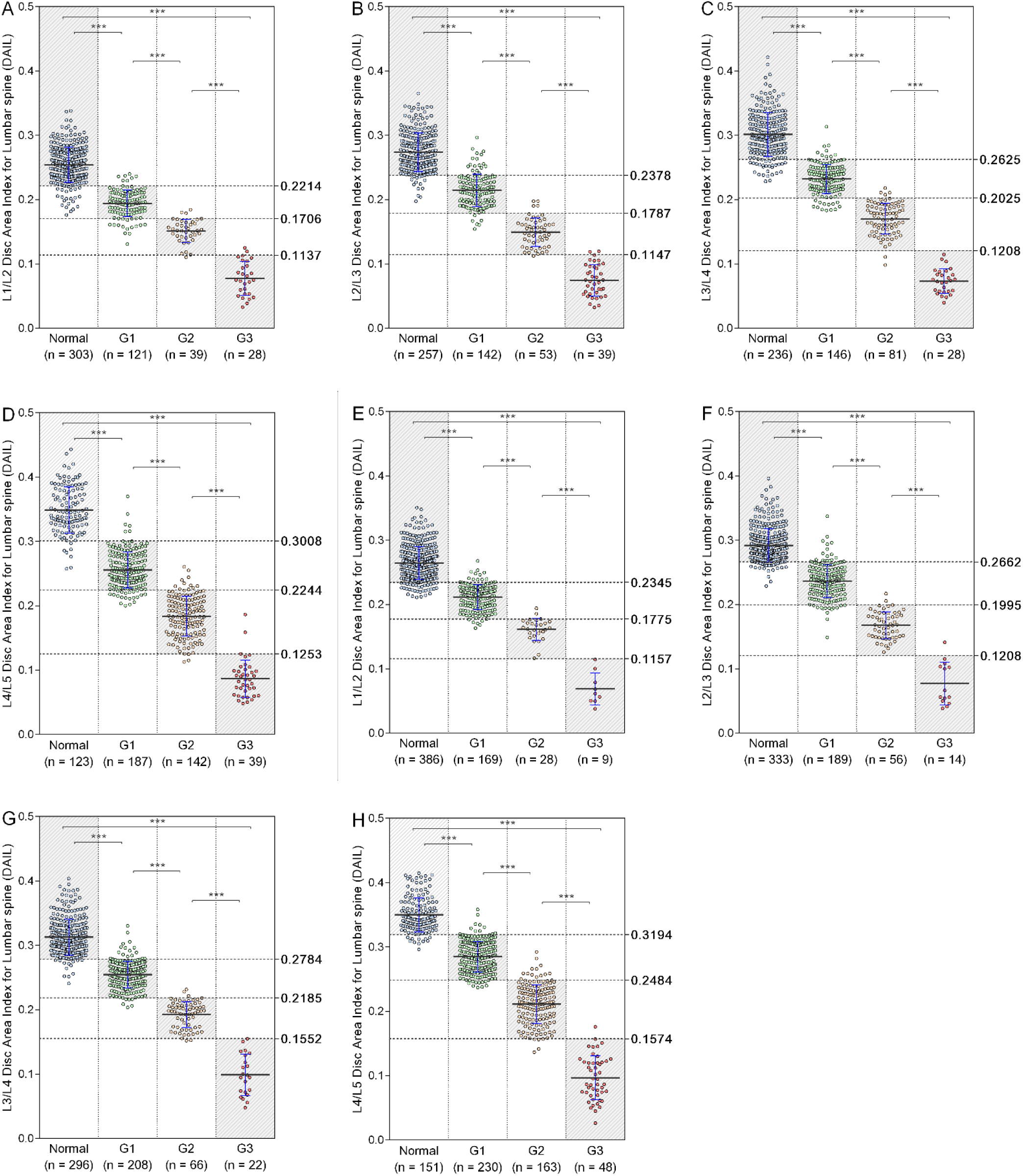
Correlations between each Individual DAIL (Disc Area Index for Lumbar Spine) and defined semi-quantitative grading at different disc Levels among women and men. A, Female L1/L2. B, Female L2/L3. C, Female L3/L4. D, Female L4/L5. E, Male L1/L2. F, Male L2/L3. G, Male L3/L4. H, Male L4/L5.

**Supplement Figure 4.**
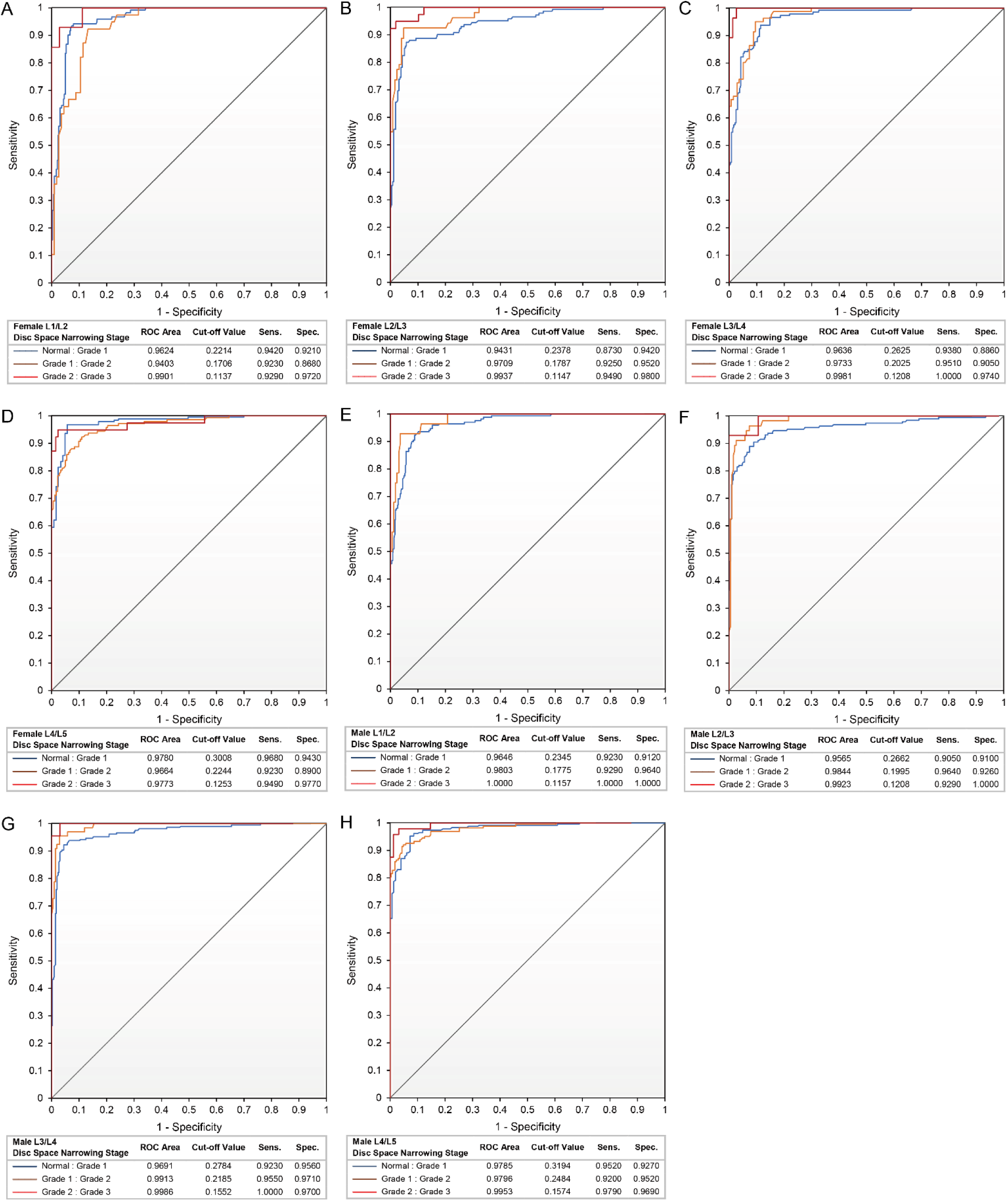
ROC analysis of the DAIL (Disc Area Index for Lumbar Spine) based classification ability for grading disc space narrowing among women and men. **A,** Female L1/L2. **B,** Female L2/L3. **C,** Female L3/L4. **D,** Female L4/L5. **E,** Male L1/L2. **F,** Male L2/L3. **G,** Male L3/L4. **H,** Male L4/L5.

**supplementary table 1.** Ha: anterior disc height; Hm: middle disc height; Hp: posterior disc height. Disc heights are presented in pixels which have adjusted that the sizes are the same for baseline and for follow-up.

| Thoracic/lumbar disc heights at baseline and year-4 follow-up among elderly females |  |  |  |  |  |  |  |  |  |
| --- | --- | --- | --- | --- | --- | --- | --- | --- | --- |
| Disc Level | Baseline |  |  | Year-4 Follow-up |  |  | 4-Years decrease (%) |  |  |
|  | Ha | Hm | Hp | Ha | Hm | Hp | Ha | Hm | Hp |
| T4/5 | 31.2 ± 9.0 | 52.3 ± 13.6 | 36.6 ± 11.9 | 30.9 ± 9.1 | 51.4 ± 12.7 | 34.7 ± 10.6 | -0.4 ± 0.18 | 0.2 ± 0.15 | 1.1 ± 0.25 |
| T5/6 | 35.3 ± 9.7 | 51.3 ± 13.2 | 35.4 ± 11.6 | 34.9 ± 9.9 | 50.4 ± 13.1 | 33.3 ± 10.4 | 0.2 ± 0.16 | 0.9 ± 0.14 | 2.1 ± 0.25 |
| T6/7 | 40.1 ± 11.4 | 53.1 ± 13.6 | 35.3 ± 10.1 | 39.1 ± 11.4 | 51.4 ± 13.3 | 33.8 ± 10.1 | 1.3 ± 0.19 | 2.4 ± 0.13 | 2.2 ± 0.21 |
| T7/8 | 44.5 ± 12.6 | 55.5 ± 14.2 | 34.9 ± 10.1 | 43.0 ± 12.2 | 53.3 ± 13.6 | 33.2 ± 9.8 | 2.0 ± 0.17 | 2.8 ± .14 | 3.0 ± 0.19 |
| T8/9 | 47.8 ± 13.8 | 58.2 ± 14.8 | 34.1 ± 9.6 | 46.5 ± 13.7 | 56.0 ± 14.2 | 32.6 ± 10.0 | 1.8 ± 0.15 | 2.8 ± .13 | 2.8 ± 0.21 |
| T9/10 | 53.3 ± 15.0 | 62.4 ± 15.4 | 35.8 ± 10.3 | 51.7 ± 13.7 | 60.1 ± 14.5 | 34.3 ± 9.9 | 1.4 ± 0.15 | 2.7 ± .12 | 2.1 ± 0.20 |
| T10/11 | 60.68 ± 6.1 | 70.7 ± 16.6 | 42.3 ± 11.7 | 58.8 ± 15.5 | 68.1 ± 16.1 | 40.4 ± 11.4 | 2.0 ± 0.14 | 3.1 ± .11 | 3.3 ± 0.16 |
| T11/12 | 69.8 ± 18.2 | 82.1 ± 19.0 | 51.5 ± 13.9 | 67.2 ± 17.2 | 78.9 ± 18.5 | 48.9 ± 13.3 | 2.7 ± 0.14 | 3.3 ± .11 | 3.7 ± 0.16 |
| Thoracic discs (mean) | 48.1 ± 18.2 | 60.6 ± 18.1 | 38.3 ± 12.6 | 46.7 ± 17.4 | 58.6 ± 17.3 | 36.4 ± 12.0 | 1.4 ± 0.16 | 2.3 ± .13 | 2.5 ± 0.21 |
| T12/L1 | 80.7 ± 21.7 | 95.4 ± 22.9 | 55.1 ± 15.2 | 77.2 ± 20.9 | 92.5 ± 22.9 | 52.5 ± 15.2 | 3.6 ± 0.11 | 2.7 ± .10 | 3.7 ± 0.16 |
| L1/2 | 97.2 ± 28.1 | 107.7 ± 26.6 | 63.2 ± 18.6 | 92.5 ± 26.5 | 104.6 ± 26.8 | 59.9 ± 18.4 | 4.0 ± 0.11 | 2.7 ± .10 | 4.3 ± 0.16 |
| L2/3 | 114.4 ± 33.6 | 116.7 ± 29.9 | 69.1 ± 20.5 | 109.1 ± 32.2 | 113.5 ± 31.5 | 65.0 ± 20.0 | 3.9 ± 0.12 | 2.7 ± 0.11 | 5.1 ± 0.15 |
| L3/4 | 134.3 ± 37.6 | 124.9 ± 32.4 | 75.4 ± 23.7 | 128.4 ± 36.9 | 121.4 ± 33.5 | 71.4 ± 23.0 | 3.6 ± 0.13 | 2.8 ± 0.11 | 4.7 ± 0.14 |
| L4/5 | 145.5 ± 43.3 | 127.4 ± 33.9 | 88.9 ± 28.2 | 138.7 ± 42.7 | 123.0 ± 35.7 | 84.3 ± 29.1 | 4.1 ± 0.14 | 3.5 ± 0.12 | 4.8 ± 0.15 |
| Lumbar discs (mean) | 114.8 ± 41.2 | 114.4 ± 31.7 | 70.1 ± 24.3 | 109.6 ± 39.8 | 111.0 ± 32.4 | 66.4 ± 24.0 | 3.9 ± 0.12 | 2.9 ± 0.11 | 4.5 ± 0.15 |

**supplementary table 2.** Ha: anterior disc height; Hm: middle disc height; Hp: posterior disc height. Disc heights are presented in pixels which have adjusted that the sizes are the same for baseline and for follow-up.

| Thoracic/lumbar disc heights at baseline and year-4 follow-up among elderly males |  |  |  |  |  |  |  |  |  |
| --- | --- | --- | --- | --- | --- | --- | --- | --- | --- |
| Male Disc Level | Baseline |  |  | Year-4 Follow-up |  |  | 4-Years decrease (%) |  |  |
|  | Ha | Hm | Hp | Ha | Hm | Hp | Ha | Hm | Hp |
| T4/5 | 18.3 ± 4.1 | 34.4 ± 5.2 | 22.3 ± 6.0 | 17.9 ± 4.2 | 33.7 ± 5.7 | 21.5 ± 6.8 | 0.6 ± 0.20 | 1.2 ± 0.15 | 0.4 ± 0.30 |
| T5/6 | 21.8 ± 4.7 | 34.1 ± 5.0 | 21.9 ± 5.6 | 21.2 ± 5.0 | 33.3 ± 5.3 | 21.2 ± 5.7 | 0.8 ± 0.20 | 1.6 ± 0.14 | 0.5 ± 0.25 |
| T6/7 | 25.7 ± 5.5 | 35.1 ± 5.5 | 22.1 ± 5.4 | 25.0 ± 5.5 | 34.2 ± 5.9 | 21.3 ± 5.6 | 1.1 ± 0.19 | 1.7 ± 0.15 | 0.8 ± 0.24 |
| T7/8 | 29.1 ± 5.9 | 37.2 ± 5.4 | 23.0 ± 5.6 | 28.1 ± 5.8 | 36.0 ± 6.2 | 22.0 ± 5.9 | 2.3 ± 0.16 | 2.7 ± 0.14 | 1.7 ± 0.26 |
| T8/9 | 32.0 ± 6.1 | 39.2 ± 6.0 | 23.2 ± 5.5 | 31.2 ± 6.4 | 38.0 ± 6.4 | 22.2 ± 5.6 | 1.7 ± 0.15 | 2.2 ± 0.15 | 1.5 ± 0.24 |
| T9/10 | 35.0 ± 6.7 | 42.7 ± 6.3 | 24.1 ± 5.8 | 33.7 ± 6.7 | 41.2 ± 6.7 | 23.0 ± 5.8 | 2.4 ± 0.17 | 3.0 ± 0.13 | 1.6 ± 0.25 |
| T10/11 | 38.8 ± 7.2 | 48.1 ± 7.1 | 28.6 ± 7.1 | 37.6 ± 7.0 | 46.4 ± 7.3 | 27.4 ± 6.7 | 2.2 ± 0.15 | 2.9 ± 0.12 | 1.3 ± 0.25 |
| T11/12 | 44.8 ± 8.4 | 55.3 ± 8.0 | 34.5 ± 7.4 | 43.0 ± 8.6 | 53.1 ± 8.5 | 33.2 ± 7.4 | 2.7 ± 0.19 | 3.1 ± 0.14 | 1.7 ± 0.21 |
| Thoracic discs (mean) | 30.9 ± 10.4 | 40.9 ± 9.4 | 25.1 ± 7.4 | 29.9 ± 10.1 | 39.6 ± 9.4 | 24.1 ± 7.4 | 1.7 ± 0.18 | 2.3 ± 0.14 | 1.2 ± 0.25 |
| T12/L1 | 52.0 ± 9.2 | 58.1 ± 8.2 | 35.3 ± 7.7 | 50.3 ± 9.0 | 56.5 ± 8.0 | 34.0 ± 8.0 | 2.5 ± 0.13 | 2.1 ± 0.11 | 2.7 ± 0.16 |
| L1/2 | 64.3 ± 11.4 | 66.9 ± 9.3 | 42.5 ± 8.6 | 62.2 ± 11.4 | 64.9 ± 9.8 | 40.9 ± 8.7 | 2.7 ± 0.12 | 2.6 ± 0.09 | 2.9 ± 0.16 |
| L2/3 | 74.1 ± 13.4 | 72.6 ± 10.9 | 46.4 ± 9.1 | 71.5 ± 13.7 | 70.4 ± 11.7 | 44.6 ± 9.4 | 3.0 ± 0.12 | 2.9 ± 0.09 | 3.1 ± 0.14 |
| L3/4 | 83.2 ± 14.5 | 76.0 ± 12.4 | 49.8 ± 10.4 | 80.0 ± 14.6 | 73.3 ± 12.7 | 47.6 ± 10.5 | 3.4 ± 0.11 | 3.3 ± 0.09 | 3.7 ± 0.13 |
| L4/5 | 88.4 ± 19.5 | 76.6 ± 15.3 | 52.3 ± 12.9 | 84.4 ± 19.4 | 73.5 ± 15.3 | 49.6 ± 12.9 | 3.8 ± 0.14 | 3.6 ± 0.10 | 4.1 ± 0.17 |
| Lumbar discs (mean) | 72.5 ± 19.2 | 69.9 ± 13.4 | 45.6 ± 11.5 | 69.8 ± 18.7 | 67.6 ± 13.3 | 43.6 ± 11.4 | 3.1 ± 0.12 | 2.9 ± 0.10 | 3.3 ± 0.15 |

